# Proximity sensors reveal social information transfer in maternity colonies of Common noctule bats

**DOI:** 10.1101/421974

**Authors:** Simon Ripperger, Linus Günther, Hanna Wieser, Niklas Duda, Martin Hierold, Björn Cassens, Rüdiger Kapitza, Alexander Kölpin, Frieder Mayer

## Abstract

1. Bats are a highly gregarious taxon suggesting that social information should be readily available for making decision. Social information transfer in maternity colonies might be a particularly efficient mechanism for naïve pups to acquire information on resources from informed adults. However, such behaviour is difficult to study in the wild, in particular in elusive and small-bodied animals such as bats.
2. The goal of this study was to investigate the role of social information in acquiring access to two types of resources, which are crucial in the life of a juvenile bat: suitable roosting sites and fruitful feeding grounds. We hypothesized that fledging offspring will make use of social information by following informed members of the social groups to unknown roosts or foraging sites.
3. In the present study we applied for the first time the newly developed miniaturized proximity sensor system ‘BATS’, a fully automated system for documenting associations among individual bats both while roosting and while on the wing. We quantified associations among juveniles and other group member while switching roosts and during foraging.
4. We found clear evidence for information transfer while switching roosts, mainly among juveniles and their genetically identified mothers. Anecdotal observations suggest intentional guidance behaviour by mothers, indicated by repeated commuting flights among the pup and the target roost. Infrequent, short meetings with colony members other than the mother indicate local enhancement at foraging sites, but no intentional information transfer.
5. Our study illustrates how advances in technology enable researchers to solve long-standing puzzles. Miniaturized proximity sensors facilitate the automated collection of continuous data sets and represent an ideal tool to gain novel insights into the sociobiology of elusive and small-bodied species.

## Introduction

The early development is a critical phase for an animal since it paves the way for later life by affecting survival rate and overall fitness. It has been shown for several species that limited access to food resources within the first weeks after birth has a negative impact on reproductive success (Lindström 1999; Lummaa & Clutton-Brock 2002). However, a central question is: how do offspring get access to resources in early life once they start to become independent? Behaviours related to foraging might be genetically predefined, which is the case in many invertebrates but also in vertebrate taxa (van Schaik *et al.* 2017). Over time, ‘personal information’ which is acquired by direct interaction with the environment (Dall *et al.* 2005) or individual learning by trial and error augment an individual’s capabilities (van Schaik *et al.* 2017). If animals are born and raised in presence of their parents or other conspecifics, information may be socially acquired (Galef Jr & Giraldeau 2001). Adopting information from group members may be more efficient and less costly than individual learning and contributes to an individual’s behavioural flexibility under changing conditions (Dall *et al.* 2005; van Schaik *et al.* 2017).

Information obtained within colonies or groups of conspecifics enables better decisions in various contexts such as predator avoidance, reduction of parasitism, habitat choice and foraging (Evans, Votier & Dall 2016). The mechanisms of acquiring social information may vary widely in complexity. Individuals may use ‘inadvertent social information’, which is generated by social cues of conspecifics, i.e., eating may inform about the location of food or fleeing about the presence of a predator (Dall *et al.* 2005). Such public information is created non-deliberately and in group-living animals it may be difficult to hide certain information, e.g., on foraging success. This might particularly apply for breeding colonies, where parents must return to their young and may inadvertently inform others on foraging success via, e.g., time of arrival or fatness (Evans, Votier & Dall 2016). If information is provided deliberately, ‘evolved signals’ are used to actively exchange information (Dall *et al.* 2005). Black-capped chickadees, e.g., broadcast alarm calls which contain information on the presence and even the size of a predator (Templeton, Greene & Davis 2005) and honey bees use the waggle dance to inform conspecifics on the location of a food source (Leadbeater & Chittka 2007). The ready availability of information at communal roosts gave rise to the Information Center Hypothesis (ICH), which states that such assemblages primarily evolved for the efficient exploitation of unevenly distributed food sources (Ward & Zahavi 1973). According to the ICH colony members must assess the success of returning foragers, which are later followed to food patches. However, information can also be generated right at the resource, a mechanism termed ‘local enhancement’. Black-browed albatrosses, e.g., indirectly detect food patches by approaching aggregations of foraging predators over sea (Grünbaum & Veit 2003). Both advertent and inadvertent information can be used by juveniles to find food. Juvenile rats prefer feeding sites where adults are present and scent marks and trails of adults cause juveniles to explore such sites (reviewed by Galef Jr and Giraldeau (2001)). Juveniles, since they are naive, seem to particularly rely on acquiring social information from more experienced individuals of the group like their parents (termed ‘vertical transmission’) or other adults (‘oblique transmission’) (van Schaik *et al.* 2017).

Bats are an ideal taxon to study the mechanisms of social information use in groups since the vast majority of species is gregarious and long-lived (Wilkinson & Boughman 1999; Kerth 2008; Smith, Lacey & Hayes 2017). However, there is surprisingly little known on whether and how juveniles benefit from social information provided by group members. An interesting case of information transfer across generations is reported for greater sac-winged bats. Here, vocal development of pups is influenced by imitation of territorial songs of harem males and leads to a group signature which is independent of relatedness (Knörnschild *et al.* 2010; Eckenweber & Knörnschild 2013). When it comes to learning where to roost and where to forage, knowledge on juveniles becomes scarce and existing literature focusses on horizontal information transfer among adult peers. Empirical studies in the wild demonstrated that several insectivorous species are attracted to feeding buzzes by conspecifics which may be used as a signal of foraging success (Gillam 2007; Dechmann *et al.* 2009; Cvikel *et al.* 2015). Female greater spear-nosed bats coordinate their foraging bouts at the day roost by screech calls (Wilkinson & Boughman 1998). However, neither local enhancement nor recruitment at the roost has been demonstrated in foraging juveniles, so far. Spatial association among home ranges of mothers and offspring in at least three species and simultaneous feeding of mother-pup pairs in vampire bats suggest that vertical information transfer, possibly in form of following behaviour, might provide juveniles with insights on where to forage, but this mechanism has yet has to be demonstrated (Wilkinson 1995; Schnitzler, Moss & Denzinger 2003).

Similarly little is known on how juveniles learn about the location of suitable roosts and the few existing studies only involved adults. In Common noctule bats local enhancement by inadvertent acoustic cues significantly reduces the time required to locate a roost both in captive and in wild experiments (Ruczyński, Kalko & Siemers 2007; Furmankiewicz *et al.* 2011). Kerth and Reckardt (2003) tracked nightly roost switching behaviour in Bechstein’s bats and assumed that recruitment of naïve by informed individuals already started at the day roost, but could not unequivocally prove it. An exclusion experiment by Wilkinson (1992) demonstrated that juvenile bats are able of relocating at a new roost with their mothers and the author concluded that following behaviour is the only plausible explanation. One of the still standing mysteries in bat ecology is how juveniles of temperate species locate swarming or hibernation sites, which are often long distances from where they are born and reared. It has been hypothesized that mothers guide their offspring (Sachteleben 1991), but so far nobody was able to track such guidance behaviour. These examples emphasize that there are plenty of indications on following and guidance behaviour in juvenile bats, but so far technological limitations prevented the final proof.

While emerging technologies have revolutionized the field of bio-logging and in turn our understanding of behaviour of wild animals during the past decades, studies on small-bodied vertebrates still lag behind due to the scarcity of fully automated lightweight devices (Kays *et al.* 2015). Proximity loggers represent a powerful tool for the study of information transfer (Rutz *et al.* 2015; St Clair *et al.* 2015), but studies making use of such devices are generally rare and the loggers used in the aforementioned studies are by far too heavy for tagging medium-sized bats. Smaller tag versions of acceptable weight, however, show dramatically reduced runtimes of less than 24 h (Levin *et al.* 2015). In the present study we used the newly developed miniaturized proximity sensor system ‘BATS’, a fully automated system for documenting associations among individuals at a tag weight of one to two gram and runtimes of at least one to two weeks (Duda, Weigel & Koelpin 2018). Our developments enable us to study interactions among tagged bats both while roosting and while on the wing. Here we report on the first extensive study to apply our system and proximity sensors to free-ranging bats, in general.

The goal of this study was to investigate the use of social information in acquiring access to two types of resources, which are crucial in the life of a juvenile bat: suitable roosting sites and fruitful feeding grounds. We hypothesized that fledging offspring will make use of social information by following either the mother or other informed members of the social groups to unknown roosts or foraging sites. If juveniles use social information when switching roosts, we expect that the successfully switching juvenile will be associated with at least one individual of the group shortly before and shortly after leaving the current roost and shortly before and shortly after arriving at the new roost. If social information is used for finding foraging grounds, we would expect juveniles to be associated with at least one roosting partner shortly before and shortly after starting the bout, during several minutes after starting while commuting to the foraging ground, and possibly, but not necessarily when returning to the roost. The BATS-tracking system enabled us to classify and quantify the aforementioned events.

## 2. Materials and methods

### 2.1 Field site and study species

This study was conducted in “Königsheide Forst”, a mixed forest in Berlin, Germany, from June to August 2016 and 2017, respectively. The study site comprises ample of roosting opportunities for bats such as natural tree holes and roughly 130 bat boxes. During this time of the year females of the common noctule bat (*Nyctalus noctula*) form temporary groups, so called maternity colonies, to jointly give birth and rear their young. Mothers give birth to one or two offspring and individuals of a maternity colony frequently switch roosts, but usually stay within the area of the nursing colony. Moving among roosts may involve a change in group composition. However, strong, non-random inter-individual bonds have been observed in captive studies as well as a certain degree of maternal care such as allogrooming of offspring (Kleiman 1969; Kozhurina 1993).

The ideal opportunity to observe information transfer in maternity colonies should be the moment when the offspring start to fledge in order to track their behaviour during the first nights of independent flight. Therefore, we daily monitored the bat boxes, including checks after sunset when adults and already flying juvenile had emerged from the roost. We aimed at tagging the majority of a social group including juveniles, which have started fledging only recently or which not fledged at all, yet. We therefore prepared to capture on the following day when around a third of the offspring were still inside the roost while the rest of colony (including already fledged youngsters) was foraging.

### 2.2 Sample collection, molecular analysis and identification of mother pup pairs

In 2016 we captured a social group from a single bat box while in 2017 bats were caught from two different bat boxes which were roughly 300m apart. Bats were kept in cotton cloth bags until they were weighted, sexed and the forearm was measured using a calliper. If the epiphyseal gaps were closed and the phalangeal–metacarpal joints were knobby, individuals were considered adult (Brunet-Rossinni & Wilkinson 2009). We collected tissue samples with a biopsy punch (Ø 4 mm, Stiefel Laboratorium GmbH, Offenbach, Germany) and preserved them in 80% ethanol. In the lab we used the salt–chloroform procedure (Miller, Dykes & Polesky 1988) modified by Heckel *et al.* (1999) for DNA isolation.

We used the DNA Analyser 4300 and the SAGA^GT^ allele scoring software (both: LI–COR Biosciences, Lincoln, NE, USA) to genotype a total of 75 individuals (n□= 33 adult females, n□l=□42 juveniles) at 9 polymorphic microsatellite loci. We used the loci P11, P217, P219 and P223 which were isolated from the focus species *Nyctalus noctula* (Mayer, Schlötterer & Tautz 2000). Nleis3 and Nleis4 were isolated from the closely related *Nyctalus leisleri* (Boston, Montgomery & Prodöhl 2009) and G6-Mluc, G31-Mluc, H23-Mluc and H29-Mluc have originally been isolated from *Myotis myotis* (Castella & Ruedi 2000), but were subsequently modified for cross-species utility in vespertilionid species (Jan *et al.* 2012). To calculate allele frequencies all adult individuals from both years (n=33) were used. All individuals were genotyped at least at eight loci, and genotypes were 99.7% complete. See Table S1 in supporting information for allele numbers per locus, results of Hardy–Weinberg tests, null allele frequencies, and non-exclusion probabilities for the nine microsatellite markers.

Parentage analyses were performed with CERVUS v. 3.0 (Kalinowski, Taper & Marshall 2007) separately for the social groups caught in 2016 and 2017, respectively, since our objective was to identify mother-pup pairs within year, not across years. The 2016 data set comprised 20 juveniles and 13 adult females (candidate mothers), while 22 juveniles and 24 candidate mothers were used for 2017. Four of the 24 adult females in 2017 were recaptures that were already caught in 2016 as juvenile (n=1) and adults (n=3).

Simulations were run with 100□000 cycles, a proportion of 80% sampled candidate mothers, an estimated genotyping error of 2%, and for two confidence levels (80% and 95%). One mismatch per mother–offspring dyad was accepted to account for genotyping errors or mutations. A mother could be assigned to 40 (2016: n=18; 2017: n=22) of the 42 analysed juveniles. Thirty-three mother-pup pairs were assigned at 95 % confidence with no mismatch, six at 95 % confidence with one mismatch and only one with 80 % confidence and one mismatch.

### 2.3 Automated encounter detection among tagged bats

Our team developed a tracking system for direct encounter detection, which bases on wireless sensor network technology for field strength related distance estimation between individuals. The system is fully automated including remote data download and does not require recapturing tagged animals thus reducing disturbance of the animals to a minimum. The centrepiece of the tracking hardware is the animal-borne mobile node, in the following referred to as ‘proximity sensor’. Once deployed, a wake-up receiver on the proximity sensor permanently scans its surroundings for signals of other proximity sensors, which are constantly broadcasted every two seconds. This operation mode is independent of any further infrastructure. Whenever one or more tracking sensors are within reception range of ca. 10 m maximum distance (Ripperger *et al.* 2016), a so called ‘meeting’ is created. As soon as no signal has been received by the respective meeting partner for five sending intervals (corresponding to 10 seconds), the meeting is closed and stored to on-board memory along with the ID of the meeting partner, a timestamp, total meeting duration, and a maximum signal strength indicator (RSSI) of the meeting. The signal, which is broadcasted every 2 s, is simultaneously used as an indicator of presence at a site of interest, e.g. a roost, when the signal is received by a stationary node, in the following referred to as ‘base stations’. We positioned base stations near potential roosts to detect presence of individual tagged bats inside a particular bat box or tree hole and we therefore termed a bat signal which are picked up by base stations ‘presence signal’. Base stations also provide remote data download, while all downloaded data is locally stored and can be accessed by the user. In 2016 the system could operate a maximum of 30 IDs at a time, while in 2017 the maximum number of observable individuals has been increased to 60. In the following we give a brief overview of the hardware components and the functionality of the system. For an elaborate, in-depth description of the software see Cassens *et al.* (2017) and for hardware see Duda, Weigel and Koelpin (2018).

#### 2.3.1 Proximity sensors

We used a refined version of miniaturized proximity sensors, which has been described and tested in free-ranging bats first in Ripperger *et al.* (2016). The proximity sensor comprises a System-on-Chip (SoC) for communication control and on-board data processing, a transceiver which enables communication in the 868MHz frequency band with other proximity sensors or base stations and a wake-up-receiver which activates full system functionality from an energy-saving low-power mode whenever communication partners are in range. A lithium-polymer battery powers the mobile node. We built two versions of the proximity sensor that differ in weight since adult females and offspring of noctule bats varied considerably in body weight. The low-weight version for tagging offspring was equipped with a 15 mAh battery and was housed in the fingertip of a nitrile lab glove. The heavier version for adult females was equipped with either two 15 mAh battery of a single 24mAh batteries and housed in a 3D-printed plastic case ensuring longer runtime. The different proximity sensor versions resulted in a total weight of 1.1 to 1.9 g including battery and housing.

#### 2.3.2 Base stations and data access

The base station contains a receiver for the reception of presence signals and transmitted data. Presence signals and downloaded data are stored by a Raspberry Pi (Raspberry PI Foundation, Cambridge, UK) to a SD card along with the ID of the transmitting proximity sensor and the receiving base station, respectively, and a timestamp which is provided by a GPS unit. At the same time the Raspberry Pi hosts a WiFi hotspot allowing the user remote data access. The data is then stored in a MySQL database.

### 2.4 Tagging and data collection

On July 15^th^ 2016 we tagged in total 26 individuals, 10 juveniles and 16 adult females, from a single bat box. On July 18^th^ 2017 we tagged in total 34 individuals, 19 juveniles and 15 adult females from two bat boxes. This adds up to a total of 60 tracked bats, 29 of which were juveniles and 31 were adult females. According to individual body weight we used different versions of the proximity sensors. Body weight ranged from 17g to 25g for juveniles and averaged at 21.26g +/− 2.04, while adult body weight ranged from 23.5g to 35g at an average of 27.73g +/− 2.27. Individual tag-to-body weight ratios ranged from 4.4% to 7% for juveniles and from 4.2% to 8% in adults, which is well within the recommendations for short-term biologging studies in bats (Amelon *et al.* 2009; O’Mara, Wikelski & Dechmann 2014). Proximity sensors were glued to the fur on the back of the bats using surgical cement (Perma-Type, Plainville, CT, USA) and drops off when the cement loses its tackiness.

Data collected during the first night after the tagging event was discarded to account for potential behavioural changes right after tagging and actual data collection started the night after in order to allow the bats to get used to the tag. In 2016 data collection lasted until July 28^th^ (12 days) and in 2017 until August 8^th^ (20 days). We installed three respectively five base stations in 2016 and 2017 at day roosts to download data and to receive presence signals for individual bats. Whenever bats switched to unknown roosts we used a handheld 868 MHz panel antenna (HSP-868C, WiMo, Herxheim, Germany) connected to a base station to localize the unknown roost and relocate base station.

### 2.5 Analysis of tracking data

We used the library RMySQL in R (James & DebRoy 2012) to access the data, which were managed in HeidiSQL, a Windows client for MariaDB. In a first step we plotted and visually explored presence signals and meetings received at base stations to define foraging bouts and roost switching events, respectively, for all individual juveniles (see Fig. 2a-c for examples). To evaluate potential information transfer we queried the meeting database for events, which matched the timestamp of foraging bouts or roost switches, respectively. If information transfer would play a role during foraging or roost switching, we would expect to find meetings among offspring and other group members associated with these events. In detail, we proceeded as follows.

**Figure 1:**
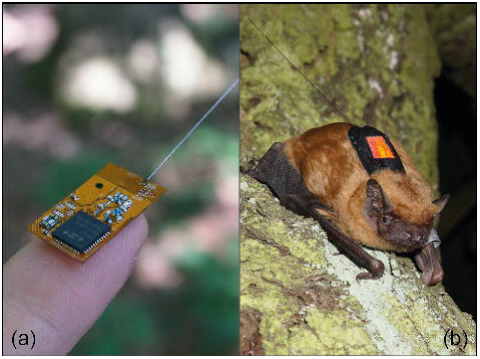
Unpackaged proximity sensor (a) and tagged adult Common noctule bat (*Nyctalus noctula*) ready for take-off (b).

**Figure 2:**
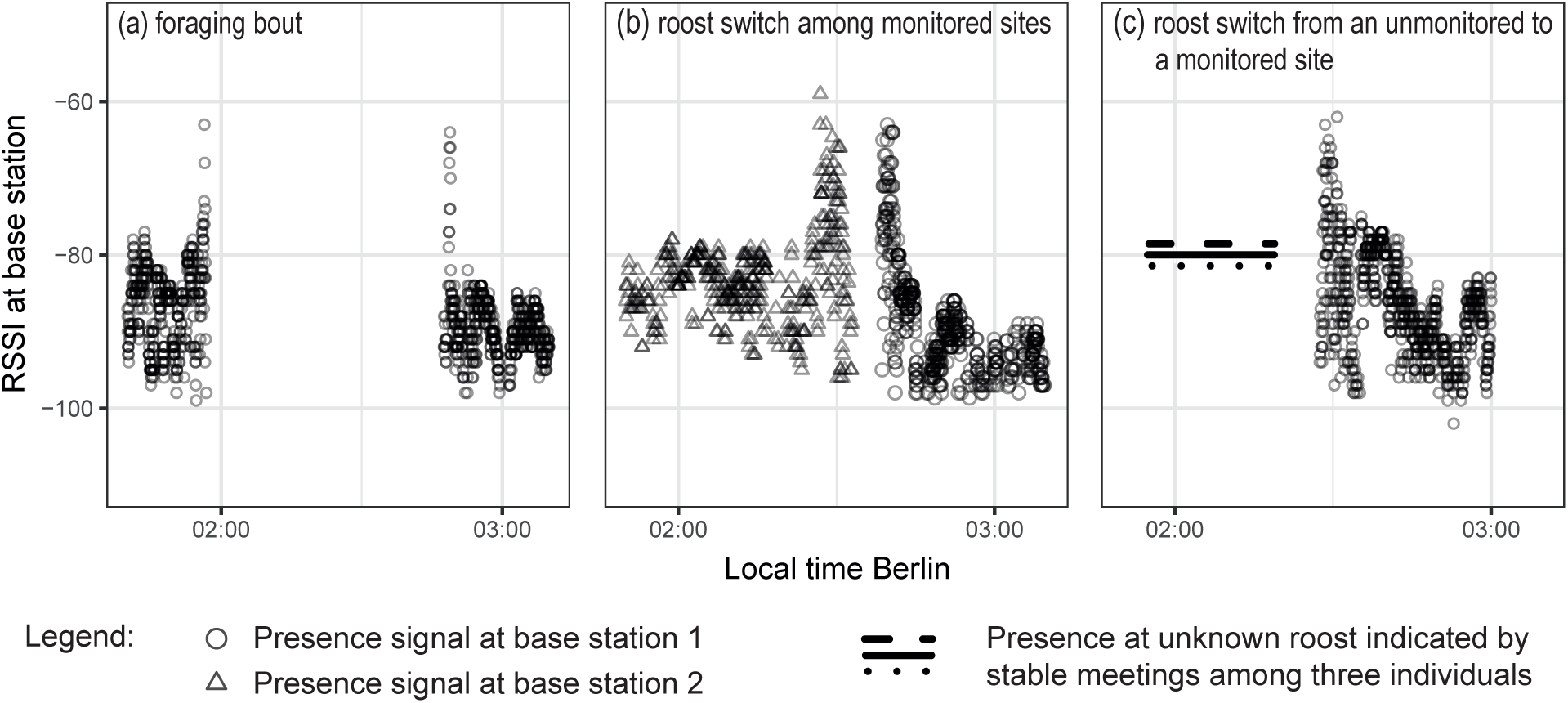
Visual representation of foraging bouts and roost switches based on presence signals at bat boxes (base stations) and meeting data. (a) A foraging bout is characterized by an interrupt of the presence signals of an individual bat which are received by a base station at a specific roost. Usually, variation of the received signal strength indicator (RSSI) increases when a bat is leaving a roost compared to when it is roosting (notice the pronounced spike upon departure and return). (b) A roost switch among two monitored sites is displayed. The presence signals interrupt at base station two while the strong variation in RSSI indicates that the bat is flying. Presence signals are then received by base station 1. (c) A roost switch occurs among an unmonitored to a monitored site. Roosting at the unmonitored site is indicated by long-lasting stable meetings among three bat individuals. Meetings interrupt when a bat individual leaves the unmonitored site followed by signal beacons being received by the base station at the monitored site.

#### 2.5.1 Evaluation of information transfer during roost switching

We defined a roost switch as an event during which an individual changes its roost and potentially its roosting partners without prolonged absence times which may indicate foraging. A roost switch can be detected if a bat switches between two roosts which are both equipped with a base station receiving presence signals (Fig. 2b). If at least one roost is equipped with a base station, presence signals can be used to determine departure time or arrival, respectively. If the unmonitored roost is occupied by other tagged bats (indicated by reciprocal, stable meetings) we can at least unequivocally classify this event as a roost switch (Fig. 2c). However, we cannot determine the time of arrival respectively departure at the unmonitored roost because bats may leave jointly. If juveniles use social information when switching roosts, we expect that the switching juvenile will be associated with at least one individual of the group shortly before and shortly after leaving the current roost or arriving at the new roost. To this end we define the moment of departing from or arriving at a monitored roost, respectively, when the steady reception of signal beacons at a base station gets cut off or starts. We subsequently queried the meeting database for meetings which are active or which started within 60s before and within 60s after the moment of leaving or arriving at a roost.

#### 2.5.2 Evaluation of information transfer during foraging bouts

We defined a foraging bout as an event where an individual starts from a known roost, returns to the same roost and does not visit other monitored roosts or roosts with tagged bats (indicated by stable, lasting meetings) in between (Fig. 2a). We chose these strict rules to ensure that the events we are looking at relate to foraging and do not overlap with roost switching events. If social information would play a role in locating foraging grounds we would expect a juvenile to associate with at least one roosting partner upon starting the bout, during several minutes after departure while commuting to the foraging ground, and possibly, but not necessarily when returning to the roost. As described above we equally defined the start and the end of the foraging bout as the end and the start of the steady reception of the presence signal, respectively. We then queried all meetings which were ongoing or started within 60s before and within 60s after starting a foraging bout and returning, respectively. In addition, we queried all meetings which originated during the entire foraging bout.

#### 2.5.3 Statistical testing

We used a Mantel-Test to test whether social information used by juveniles is obtained by their mothers in the first place or by any roosting partner. To this end we created a binary matrix containing “1” for dyads which have been associated while roost switching and “0” for dyads which have never been observed switching communally. Accordingly, foraging associations were transformed into a binary martrix. For testing the effect of maternity we created a second binary correlation matrix which listed the genetically determined identity of mother-pup pairs as “1”, while all other dyads were marked “0”. We tested the years 2016 and 2017 separately and ran Mantel tests in the library “ade4” version 1.7-11 in RStudio 1.1.453 using Monte-Carlo permutation tests with 9999 replicates (Dray & Dufour 2007; R Developing Core Team 2015).

## 3. Results

### 3.1 Genetic analyses

Mother and juvenile bats were caught in day roots at the time of weaning. In 24 determined mother-pup pairs, both individuals were tagged with proximity sensors (2016: n=9, all assigned at 95 % confidence with no mismatch; 2017: n=15, 12 pairs assigned at 95 % confidence with no and three pairs at 95 % confidence with one mismatch). These 24 mother-pup pairs generated the data for the following section.

### 3.2 Tracking results

In 2016 we received a total of 561,795 presence signals and 13,292 meetings from 23 individual bats and in 2017 we received 2,667,409 localization signals and 53,391 meetings from 33 individual bats. One individual in 2016 and three individuals in 2017 did not get in contact with base stations. These four individuals may have left the study area between tagging and the following night.

#### 3.2.1 Evaluation of joint roost switching events

To evaluate information transfer on roosts we screened the data set for joint departures from and joint arrivals at roosts for all tagged juveniles. In 2016 we observed ten events of seven individual juveniles being associated with another individual while switching among two roosts. In all except one event the associated bats arrived together at a new roost, even though successful switching took several approaches in two cases and temporary roosts may be used in between (Table 1, Fig. 3). In six cases both roosts have been monitored by a base station, in two cases the juveniles left a monitored roost and switched to a roost where other tagged bats have been roosting and in the remaining two cases the juveniles switched from a monitored roost to an unknown roost where no other tagged bats were present, except the one which accompanied the juvenile during switching. In all 10 cases the juvenile was in company of its identified mother and no other tagged bat.

**Figure 3:**
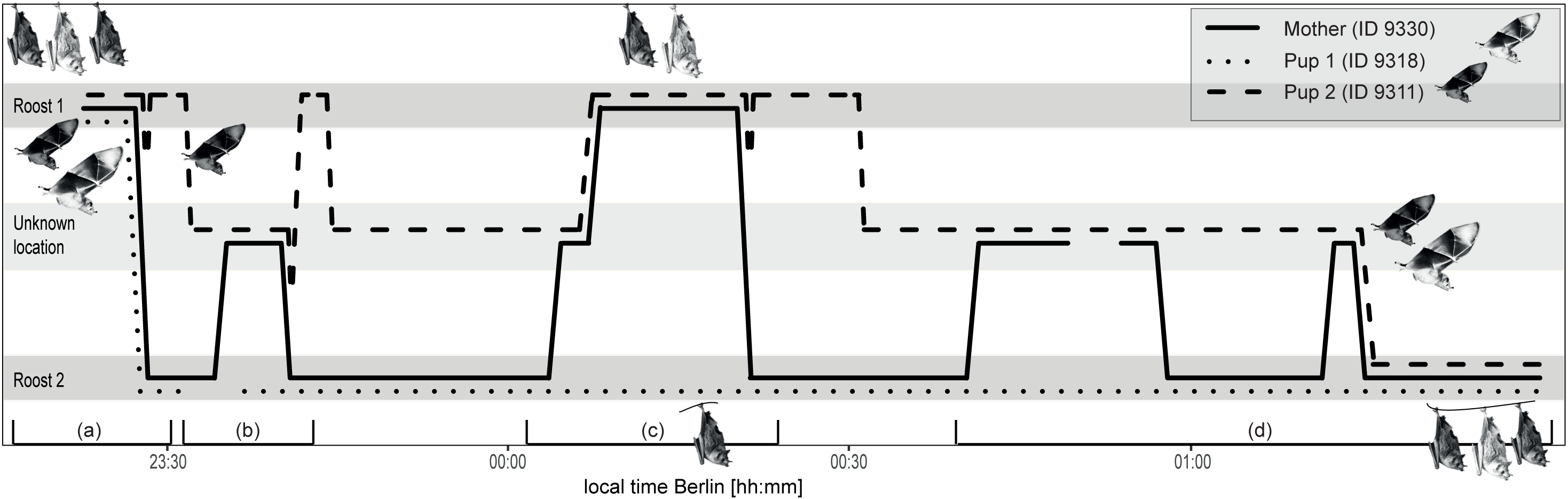
Schematic representation of a mother and its twins switching roosts: repeated commutes indicate intentional behaviour of the mother. (a) A mother and its twins jointly leave roost 1 and the mother successfully transfers to roost 2 with pup 1. The meeting to pup 2 aborts and pup 2 is flying back to roost 1. (b) Pup 2 moves solitarily from roost 1 to an unknown location where it is joined by its mother after a few minutes. Both fly in company towards roost 2, but pup 2 flies back to roost 1 while a meeting starts among the mother and pup 1 at roost 2. (c) The mother joins pup 2 in an unknown location and they jointly switch to roost 1. They jointly leave roost 1, but only the mother arrives at roost 2 starting a meeting with pup 1, while the pup 2 flies back to roost 1. (d) The mother joins pup 2 in an unknown location, around 00:50 am the meeting is interrupted for several minutes (possibly because at least one individual left), before the mother commutes twice between its two pups. Finally, around 01:15 am the mother successfully switches with pup 2 to roost 2 while a meeting is ongoing. The triad stays at roost 2 until shortly before 2 am.

**Table 1:** Summary of joint roost switching events of pups and associated partners. Switching durations were only determined when both roosts were known and equipped with base stations, while NA represents unmonitored sites or uncertain switching mode.

|  |  | partnerID | start time | end time | duration | Switching |
| --- | --- | --- | --- | --- | --- | --- |
| pupID | sex | (mother) | [date-time] | [date-time] | [hh:mm:ss] | mode |
| 9307 | f | 9338 (m) | 2016-07-21 02:59:35 | NA | NA | NA |
| 9311 | f | 9330 (m) | 2016-07-16 23:26:22 | 2016-07-17 01:15:06 | 01:48:44 | 3* |
| 9311 | f | 9330 (m) | 2016-07-20 03:18:35 | 2016-07-20 03:19:31 | 00:00:56 | 1 |
| 9318 | f | 9330 (m) | 2016-07-16 23:26:33 | 2016-07-16 23:28:31 | 00:01:58 | 1* |
| 9318 | f | 9330 (m) | 2016-07-20 04:14:25 | 2016-07-20 04:14:54 | 00:00:29 | 3 |
| 9319 | f | 9327 (m) | 2016-07-16 22:48:01 | NA | NA | NA |
| 9312 | f | 9334 (m) | 2016-07-17 01:29:14 | NA | NA | 1 |
| 9323 | m | 9340 (m) | 2016-07-16 23:58:13 | 2016-07-17 01:37:06 | 01:38:53 | 2 |
| 9323 | m | 9340 (m) | 2016-07-19 02:05:26 | NA | NA | 1 |
| 9325 | m | 9336 (m) | 2016-07-17 02:44:31 | 2016-07-17 03:55:19 | 01:10:48 | 2 |
| 9376 | f | 9383 | 2017-07-20 04:23:43 | 2017-07-20 04:23:49 | 00:00:06 | 1 |
| 9376 | f | 9383 | 2017-07-22 02:25:53 | NA | NA | 1 |
| 9370 | f | 9368 (m) | 2017-07-20 04:35:41 | NA | NA | 1 |
| 9373 | m | 9412 | 2017-07-22 02:38:13 | 2017-07-22 02:38:39 | 00:00:26 | 1 |
| 9380 | m | 9413 | 2017-07-20 04:17:52 | NA | NA | 1 |
| 9391 | f | 9385 (m) | NA | 2017-07-22 02:22:25 | NA | 1 |

In 2017 we observed six events where 5 individual juveniles switched roosts in company. Twice, the juvenile switched among two monitored roosts and four times among one monitored and an unmonitored site. Twice, the juvenile was associated with its identified mother, in four cases with another adult female.

Some juveniles switched directly among roosts. Such events took only seconds to minutes (see table xx). During other events stopover sites were used and several attempts of mothers re-associating with their young were necessary before both arrived at the new roost. Such unsuccessful tandem flights underline that the offspring was actively flying and not carried by the mother.

In both years significantly more mother-pup dyad have been observed switching roosts communally than expected by chance (Mantel tests, 9999 replicates; 2016: r = 0.88, p < 0.001; 2017: r = 0.21, p < 0.01).

#### 3.2.2 Associations during foraging bouts

In total we detected 42 foraging bouts of juveniles, which matched our definition above, conducted by 13 individuals (2016: four juveniles, eight bouts; 2017: nine juveniles, 34 bouts). Foraging bouts lasted on average 1:14:53 h with a standard deviation of 36:19 min. During 6 of these 42 foraging bouts (14 %, n = 7 individual juveniles, all 2017) we detected in total 28 short meetings, which lasted between 1 and 30 seconds (average: 7.4 s +/− 8.6). Two of these meetings occurred within less than 90 s after two co-roosting individuals left a roost; however, no further meetings have been documented during these foraging bouts. All remaining meetings originated at least several minutes after emergence from the roost. Eight times the meeting partner was another juvenile and twice an adult female. Only in one case the meeting partner was the identified mother. Accordingly, meetings among mother-pup dyads have not been observed more often than expected by chance (Mantel tests, 9999 replicates, r = 0.08, p > 0.05).

In 13 out of the 42 foraging bouts of juveniles (2016: n = two individuals; 2017: n = seven individuals) the identified mother was co-roosting before both started a bout. In all 13 cases the mother started its foraging bout considerably earlier than the juvenile (between 4:31 min and 1:26:02 h, average: 36:45 min (+/− 24:57)).

## Discussion

The study of information transfer in free-ranging bats is particularly challenging due to their small body size and their elusive, nocturnal life. We tracked bats using novel, miniaturized proximity sensors and demonstrated that juveniles use social information of group members and for finding roosts mothers seem to intentionally guide their young. However, during foraging mothers did not guide their offspring, but meetings with other colony members may reflect local enhancement at feeding grounds.

To the best of our knowledge our study shows for the first time that recruitment to a new roost starts already at the occupied roost. Furthermore, the repeated commuting flights we observed in at least two cases until the juvenile arrives at the target roost represents anecdotal evidence that at least in some cases deliberate, evolved signals rather than inadvertent social cues are used. The existence of evolved signals and the strong bias towards information transfer among mother-pup pairs suggests that the observed behaviour is best explained by kin selection. Some studies have reported on the use of social information in bats for finding suitable roosts, however, studies are scarce and the mechanisms are in parts poorly understood, in particular when it comes to naïve juveniles. Studies on a range of vespertilionid species including the focus species *N. noctula* have shown that conspecific calls enhance roost finding efficiency in captive experiments as well as in the wild (Ruczyński, Kalko & Siemers 2009; Schöner, Schöner & Kerth 2010; Furmankiewicz *et al.* 2011). These studies demonstrate that bats may eavesdrop on vocalizations to localize an occupied roost once within hearing distance. Since playbacks from varying contexts have been used we conclude that the studied bats relied on inadvertently broadcasted public information. On the contrary, Spix’s disk winged bats deliberately produce signals to facilitate group cohesion, by a remarkable call-and-response system among flying bats in search of a roost and bats occupying a roost (Chaverri, Gillam & Vonhof 2010; Chaverri, Gillam & Kunz 2012). A common theme of all abovementioned studies is that the mechanism of recruitment of conspecifics is best explained by local enhancement, i.e. the social information is acquired at the new roost, when searching bats are in hearing distance. Kerth and Reckardt (2003) were first to present experimental evidence for information transfer about roosts in bats. The authors presumed that naïve Bechstein’s bats are recruited to a novel roost already at the dayroost by experienced conspecifics, however, they could not unequivocally exclude local enhancement at the target roost. Our study finally demonstrates that this inferred mechanism does exist in roost-switching bats.

We classify the advertent information transfer from mothers to their young as a form of maternal care which has to the best of our knowledge not been observed in free-ranging bats, so far. Mammalian offspring is usually strongly dependent on maternal care for food, protection and warmth (Balshine 2012) and maternal investment in young is also wide-spread in bats (Smith, Lacey & Hayes 2017). Besides weaning maternal care has been demonstrated in form of post-weaning food provisioning (Wilkinson 1990; Geipel *et al.* 2013), grooming (Kleiman 1969; Wilkinson 1986; Kozhurina 1993) or pup guarding (Bohn, Moss & Wilkinson 2009). Carrying young in flight is also commonly observed and Jones (2000) summarizes some reports where young are possibly carried to temporary roosts or feeding grounds. However, this is the first study to document maternal guidance to roosts, which has been hypothesized as a plausible explanation for young to reach swarming and hibernation sites, but could not be confirmed, possibly due to the lack of appropriate tracking technology (Sachteleben 1991; Burns & Broders 2015; Stumpf *et al.* 2017).

Previous work on bats indicated that roosts may act as information centres where bats may obtain information on food by inadvertent cues (Ratcliffe & ter Hofstede 2005; O’Mara, Dechmann & Page 2014). A considerable part of the diet of Common noctule bats consists of insects, which fly in swarms and often over water (Gloor, Stutz & Ziswiler 1995). Such rich and patchy, but ephemeral foraging sites are required for the establishment of information centres (Ward & Zahavi 1973) and juveniles in particular might benefit from rich food patches when collecting experience on where and how to forage. However, we did not observe recruitment at the roost to feeding grounds in young noctules, which complies with most foregoing studies that showed that ‘ICH’ operates in colonial roosts, but is rarely demonstrated in breeding colonies (summarized by Evans, Votier and Dall (2016)). Our observation that juveniles start foraging bouts considerably later than their mothers suggests that juvenile noctules conduct opportunistic, explorative foraging flights. Rare and short contacts to tagged colony members other than the mother during foraging bouts suggest that local enhancement by eavesdropping on conspecifics while hunting may play a role as it has been shown for several insectivorous bat species (Gillam 2007; Dechmann *et al.* 2009; Cvikel *et al.* 2015). However, our data cannot unequivocally prove this theory since the exact context of the meetings remains unknown.

Our observations raise the following question: Why is social information transfer among mothers and their offspring context dependent? One possible explanation is that group cohesion is crucial for energy-saving social warming and prolonged lactation periods in bats require mothers to stay in contact with their young for 3 weeks to 2 months depending on the species (reviewed by Kerth (2008)). Extended weaning, which was observed in captive noctules for up to 2 months (Kleiman 1969), and the broad spectrum of insects they feed on (Gloor, Stutz & Ziswiler 1995) may in turn enable juveniles to forage opportunistically and – if available – make use of social information by local enhancement. In general, suitable roosts of high quality may be harder to find opportunistically than insect prey and information on roosts is likely to accumulate in adults, in particular in philopatric females. This should favour information transfer on roosts since failing to relocate at an occupied roost might be more costly than low foraging success, which might subsequently be balanced by extended weaning. Adverse climatic conditions may have detrimental effects on single bats (Lindström 1999) and might therefore be a strong driver of the evolution of the observed guidance behaviour, since local enhancement by vocalization at the roost (Furmankiewicz *et al.* 2011) might not be functional for long-distance localization of roosting partners.

## Conclusions

Bats are facing ideal prerequisites for social information transfer, since they are long-lived and the vast majority of species is living in group. Regarding information use in offspring Wilkinson and Boughman (1999) speculated already 20 years ago that young bats almost certainly follow adults in situations other than foraging. However, this is also how long it took to unequivocally track mother-pup pairs switching among roosting sites. Our study shows that the current revolution in tracking technology provides powerful tools to investigate behavioural ecology and sociobiology in free-ranging small bodied animals such as bats.

## Footnotes

**Authors’ contributions** SR and FM conceived the ideas and designed the sampling scheme; MH, ND, AK, BC and RK developed and tested all tracking equipment. SR, LG, FM and HW collected the data; SR and LG analysed the data; SR and FM led the writing of the manuscript. All authors contributed critically to the drafts and gave final approval for publication.

## Acknowledgements

We thank I. Waurick, S. Hayden and E. Jäger for support during molecular lab work. We are grateful to T. Teige for his help and his valuable expertise during field work. All necessary permits have been obtained from SenStadtUm (I E 222/OA-AS/G_1203) and LaGeSo (I C 113-G0008/16).

## Data accessibility

Data will be deposited on GFBio and made available upon acceptance.

## Supporting information

The following Supporting Information is available for this article online.

Table S1: Results from allele frequency calculations with CERVUS v. 3.0 (Kalinovski et al. 2007).

